# Mutations and predicted glycosylation patterns in respiratory syncytial virus isolates correlate with disease severity

**DOI:** 10.64898/2026.02.03.703626

**Authors:** Morgan L Hunte, Katherine W Herbst, Ian C Michelow, Steven M Szczepanek

## Abstract

Respiratory syncytial virus (RSV) remains an important cause of lower respiratory tract infections in young children, producing mild to life-threatening disease. Although rapid viral evolution through genetic drift is well established, the structural and functional impacts of specific pathoadaptive mutations linked to enhanced virulence are poorly defined. We investigated these relationships by isolating RSV from available nasal swabs of five hospitalized infants during the 2022–2023 winter season and conducting comparative viral genomic analysis. Severity of disease was evaluated using a validated clinical scoring system. Whole-genome sequencing followed by reference-guided assembly and structural modeling revealed distinct amino acid polymorphisms correlating with disease severity. Phylogenetic analysis placed all isolates within the RSV-A GA2.3.5 G clade. Isolates from mild moderate and severe cases clustered in A.D.1.5 and A.D.1.8 subclades. Nineteen amino acid differences were associated with clinical severity and isolates from moderate or severe cases replicated more rapidly in vitro than mild isolates. Computational glycosylation predictions indicated an increasing number of glycosylation sites in the G protein corresponding with greater disease severity. Together, these data suggest that specific pathoadaptive mutations may contribute to enhanced viral replication and severity, and are relevant for future surveillance efforts and the development of immune-based strategies targeting virulence-associated residues.

## Introduction

Respiratory syncytial virus (RSV) remains a leading global cause of lower respiratory tract infections in infants and young children (1–3). Virtually all children acquire RSV infection at least once by 2 years of age (4). Although premature neonates, immunocompromised individuals and those with congenital heart, chronic lung or neuromuscular disorders are at great risk for severe disease, most severe cases occur in previously healthy, full-term infants. Factors contributing to severe disease in healthy children remain unknown (5,6).

The intrinsic mutability of RSV, owing to its RNA-dependent RNA polymerase, allows it to explore a vast sequence space, generating variants with potential fitness advantages (7,8). While the fundamental biology of RSV remains conserved, multiple studies have reported that clinical outcomes may be influenced by specific, clade-defining mutations (9–16). However, this association is subject to debate owing to the influence of confounding factors such as patient age, prematurity, comorbidities, prior immunity, environmental factors, demographic and geographic variability, as well as heterogeneity in severity metrics and study methodologies (17–20).

Upon entering the human respiratory tract, RSV initiates a highly coordinated infection process driven by its eleven proteins encoded in a ∼15,000 base pair negative-sense RNA genome. Collectively, the concerted actions of these eleven proteins – NS1, NS2, N, P, M, SH, G, F, M2-1, M2-2, and L – enable RSV to induce the profound airway inflammation and epithelial injury frequently associated with RSV disease. There is a critical knowledge gap in linking specific RSV genotypes to clinical severity through integrated genomic, structural, and functional analyses (9,19). A barrier to further reducing the burden of RSV disease is the incomplete understanding of the viral and host determinants that drive the heterogeneity of clinical outcomes. This proof-of-concept study investigated the hypothesis that specific amino acid polymorphisms in RSV are associated with clinical disease severity by means of comparative genomics. We sequenced RSV isolates from children enrolled at Connecticut Children’s during a single season and correlated differences in the viral protein sequences with validated clinical disease severity scores.

## Methods

### Subjects and biospecimens

We selected eligible subjects with RSV who had been enrolled as controls in a single-center COVID-19 biorepository from November 2022 to March 2023 (IRB approval #20-055). Inclusion criteria for the current study were defined as subjects under 24 months of age who were hospitalized for treatment of RSV mono-infection confirmed by a commercial RT-PCR assay. Demographic data were collected via in-person surveys and clinical data were abstracted from inpatient records. The primary outcome was defined by the previously validated Clinical Disease Severity Score (CDSS) based on the subject’s most severe condition during their hospitalization (**S1 Table**) (22). The CDSS is an objective measure comprised of five parameters: 1) respiratory rate, 2) auscultation, 3) transcutaneous O_2_ saturation, 4) retraction, and 5) activity level (mainly assessed by feeding difficulty). Each parameter ranks from 0 (normal) to 3 (most abnormal), and subjects were classified as having mild (0-5), moderate (6-10), or severe (11-15) RSV (22). Mid-turbinate nasal fluid was collected at time of enrollment using BD Universal Viral Transport Collection Kits (ThermoFisher Scientific, Cat #B220529). Vials were vortexed for 20 seconds, media was divided into 500 μL aliquots, and samples were stored at −80 °C until used.

### Isolation of RNA from Mid-turbinate nasal swabs and total RNA QC

RNA was isolated from mid-turbinate nasal swab specimens using the Direct-zol™ RNA Miniprep Kit (Zymo Research) according to the manufacturer’s instructions. Total RNA was quantified and purity ratios determined for each sample using the NanoDrop 2000 spectrophotometer (ThermoFisher Scientific, Waltham, MA, USA). To further assess RNA quality, total RNA was analyzed on the Agilent TapeStation 4200 (Agilent Technologies, Santa Clara, CA, USA) using the RNA High Sensitivity assay.

### Illumina RNA Enrichment Library preparation and Sequencing

Total RNA samples were prepared for RNA enrichment using the Illumina RNA for Enrichment Sample Preparation with the Respiratory Virus Oligo Panel (RVOP) following the manufacturer’s protocol (Illumina, San Diego, CA, USA). Libraries were validated for length and adapter dimer removal using the Agilent TapeStation 4200 D1000 High Sensitivity assay (Agilent Technologies, Santa Clara, CA, USA), then quantified and normalized using the dsDNA High Sensitivity Assay for Qubit 3.0 (Life Technologies, Carlsbad, CA, USA). Sample libraries were prepared for Illumina sequencing by denaturing and diluting the libraries per manufacturer’s protocol (Illumina, San Diego, CA, USA). All samples were pooled into one sequencing pool, equally normalized, and run as one sample pool across the Illumina NovaSeq 6000 using version 1.5 chemistry. Target read depth of 2M reads was achieved per sample with paired end 100bp reads.

### Sequence analysis pipeline

Viral genomic analysis was performed using Illumina paired-end sequencing data processed through a reference-guided assembly pipeline. Raw reads underwent quality control with FastP and FastQC software, which trimmed low-quality bases (Q-score <20) and adapter sequences. To exclude non-RSV genetic material, unassembled but trimmed reads were taxonomically classified with Kraken2 software (23) against the standard Kraken Database. Only sequences assigned to Human orthopneumovirus (RSV taxon ID: 208895) were filtered and retained using SeqTK software. Filtered RSV-specific reads were aligned to the RSV-A reference genome (NCBI accession: KT992094.1) using BWA-MEM (v0.7.17) [28] with default settings. Consensus RSV genomes generated through reference-guided assembly were annotated using VIGOR4 (24).

### Sequence comparisons and analysis of point mutations

Genomes of the RSV clinical isolates were analyzed using Nextstrain’s web-based platform, Nextclade (v3.13.2) (25) to assign clades, identify mutations, and perform additional quality checks. Sequences were aligned with global RSV-A genomes from GenBank via Nextstrain’s automated pipeline. For mutation profiling, Nextstrain’s variant-calling module identified amino acid polymorphisms relative to the RSV A reference (EPI_ISL_412866). Clade classification followed Nextstrain’s nomenclature, supported by bootstrap values >70%.

### Culture of RSV from clinical isolates

A549 cells (ATCC CC-185) were seeded in DMEM (Gibco, Ref # 31966-021) + 1% GlutaMAX (Gibco, 35050-061) supplemented with 10% fetal bovine serum (Gibco, Ref# A56707-01) and 1% Anti-Anti (Gibco, Ref # 15240-062). Cells were allowed to adhere overnight. After 12 hours post-seed (approximately 95% confluent cells) were infected with each clinical isolate at an MOI of 0.01 for 2 hours. Inoculum was removed, cells were washed with PBS (Gibco, Ref# 2085290), and cell culture media was added. Plates were incubated in 5% CO_2_ at 37 °C for 24, 48, 72, or 96 hours post-infection (HPI).

### Determination of viral load

Extracellular viral load was determined using the 50% tissue culture infectious dose (TCID_50_) assay. The TCID_50_ assay was performed by infecting approximately 95% confluent A549 cell cultures cells in 96-well plates. Serial tenfold dilutions of each sample were prepared in eight replicates in infection medium. After washing the cell monolayers, 100 µL of each dilution was added per well and incubated at 37°C for 2 hours. An additional 75 µL of infection medium was then added, and plates were incubated for 4–5 days. After incubation, infection medium was aspirated then cells were fixed and stained with crystal violet. TCID_50_ was calculated using the Reed-Muench method.

### Structural Analyses

L polymerase structure predictions were obtained using homology-based modelling based on RSV RNA polymerase (PDB: 6UEN). N-linked and O-linked glycosylation predictions were performed using NetNGlyc 1.0 (30) and NetOGlyc 4.0 (31), respectively.

## Results/Discussion

This proof-of-concept study provides insights into molecular determinants of RSV virulence by integrating genomic and phenotypic analyses with measures of clinical disease severity. After excluding three subjects due to co-detection of SARS-CoV-2 (n=1), adenovirus (n=1), rhinovirus/enterovirus (n=1), we analyzed RSV isolates from five infants who had no comorbidities (**Table 1**). Nasal samples from the five subjects were collected a median of five days after start of symptoms (IQR, 4 - 6 days). Two subjects were scored as having severe RSV disease (Isolate 090, Isolate 104), one scored as moderate RSV disease (Is. 101), and two scored as mild RSV disease (Is. 081, Is. 084). One subject with severe disease was admitted to the intensive care unit. All subjects received oxygen therapy via nasal cannula.

**Table 1.** Cohort Characteristics.

| Characteristic | RSV Cohort<br>(n=5) |
| --- | --- |
| Enrollment Age (mos) | 9 (3-9) |
| Male | 4 (80%) |
| Race |  |
| White | 2 (40%) |
| Black | 1 (20%) |
| Other | 2 (40%) |
| Hispanic | 3 (60%) |
| Length of Stay (days) | 2 (1-3) |
| CDSS Category |  |
| Mild | 2 (40%) |
| Moderate | 1 (20%) |
| Severe | 2 (40%) |

Data given as median (IQR) or n (%)

Genome sequencing and comparative analysis of the five RSV isolates revealed that all belonged to RSV-A and the GA2.3.5 G clade. Isolates from mild cases clustered in the A.D clade with moderate and severe cases clustering in the A.D.1.5 and A.D.1.8 subclades (**Figure 1**). Comparative analysis of amino acid polymorphisms in RSV between mild and moderate/severe disease isolates revealed multiple non-synonymous substitutions. Amino acid differences at 19 positions were identified that distinguished all mild isolates from all moderate/severe isolates (**Table 2**). Of the 19 identified, the majority were found to be in the G protein (n=8) and the L polymerase (n=5). Others were found in the F protein (n=1), M protein (n=1), P protein (n=1), M2-1 protein (n=1), and M2-2 (n=2). Many are associated with altered physicochemical properties as shown in **Table 2**, which may contribute to differences in RSV disease severity observed in hospitalized children.

**Table 2.**
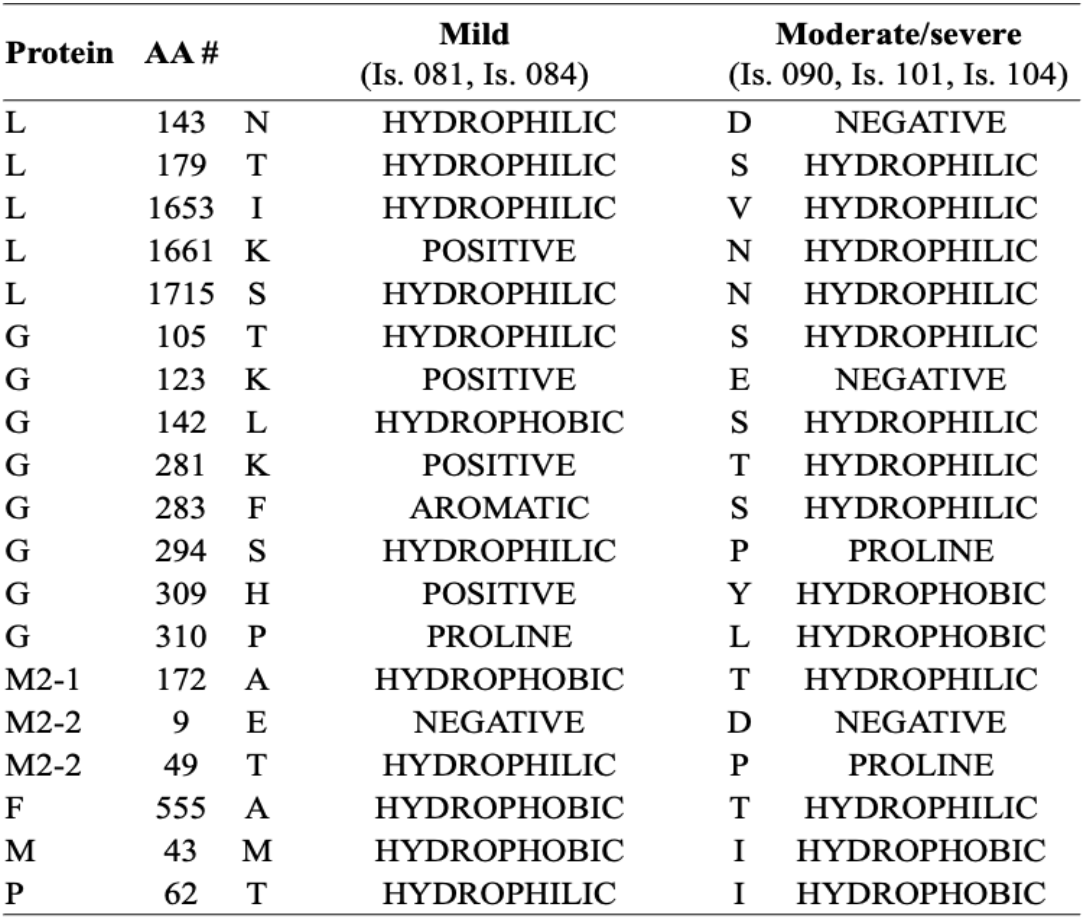
Amino Acid Differences and side chain characteristics distinguishing Mild from Moderate/Severe.

| Protein | AA # |  | Mild<br>(Is. 081, Is. 084) |  | Moderate/severe<br>(Is. 090, Is. 101, Is. 104) |
| --- | --- | --- | --- | --- | --- |
| L | 143 | N | HYDROPHILIC | D | NEGATIVE |
| L | 179 | T | HYDROPHILIC | S | HYDROPHILIC |
| L | 1653 | I | HYDROPHILIC | V | HYDROPHILIC |
| L | 1661 | K | POSITIVE | N | HYDROPHILIC |
| L | 1715 | S | HYDROPHILIC | N | HYDROPHILIC |
| G | 105 | T | HYDROPHILIC | S | HYDROPHILIC |
| G | 123 | K | POSITIVE | E | NEGATIVE |
| G | 142 | L | HYDROPHOBIC | S | HYDROPHILIC |
| G | 281 | K | POSITIVE | T | HYDROPHILIC |
| G | 283 | F | AROMATIC | S | HYDROPHILIC |
| G | 294 | S | HYDROPHILIC | P | PROLINE |
| G | 309 | H | POSITIVE | Y | HYDROPHOBIC |
| G | 310 | P | PROLINE | L | HYDROPHOBIC |
| M2-1 | 172 | A | HYDROPHOBIC | T | HYDROPHILIC |
| M2-2 | 9 | E | NEGATIVE | D | NEGATIVE |
| M2-2 | 49 | T | HYDROPHILIC | P | PROLINE |
| F | 555 | A | HYDROPHOBIC | T | HYDROPHILIC |
| M | 43 | M | HYDROPHOBIC | I | HYDROPHOBIC |
| P | 62 | T | HYDROPHILIC | I | HYDROPHOBIC |

**Figure 1.**
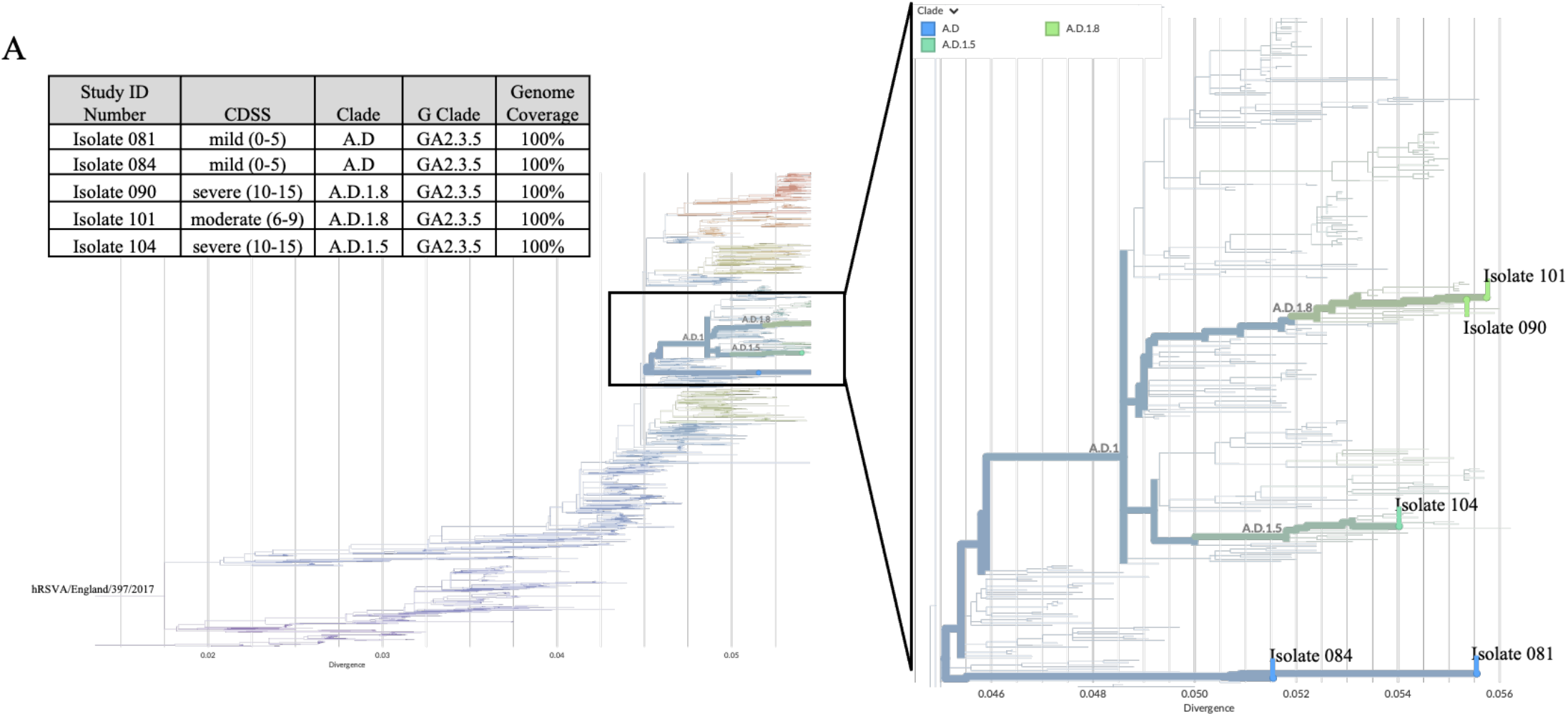
Phylogenetic analysis of RSV clinical isolates. CDSS, Clinical Disease Severity Score

In vitro growth kinetics of all RSV isolates in A549 cells demonstrated that isolates from moderate/severe cases replicated more rapidly than those from mild cases; isolates from moderate/severe cases showed significantly greater mean levels of extracellular viral titers than those from mild cases at 72 and 96-hours post-infection (two-way ANOVA with Tukey’s multiple-comparison test, p<0.05) (**Figure 2**). Importantly, the phenotypic differences in viral replication in the present study were observed in a controlled cell culture system, which reflects intrinsic viral properties because the influence of host-related confounding variables was removed.

**Figure 2.**
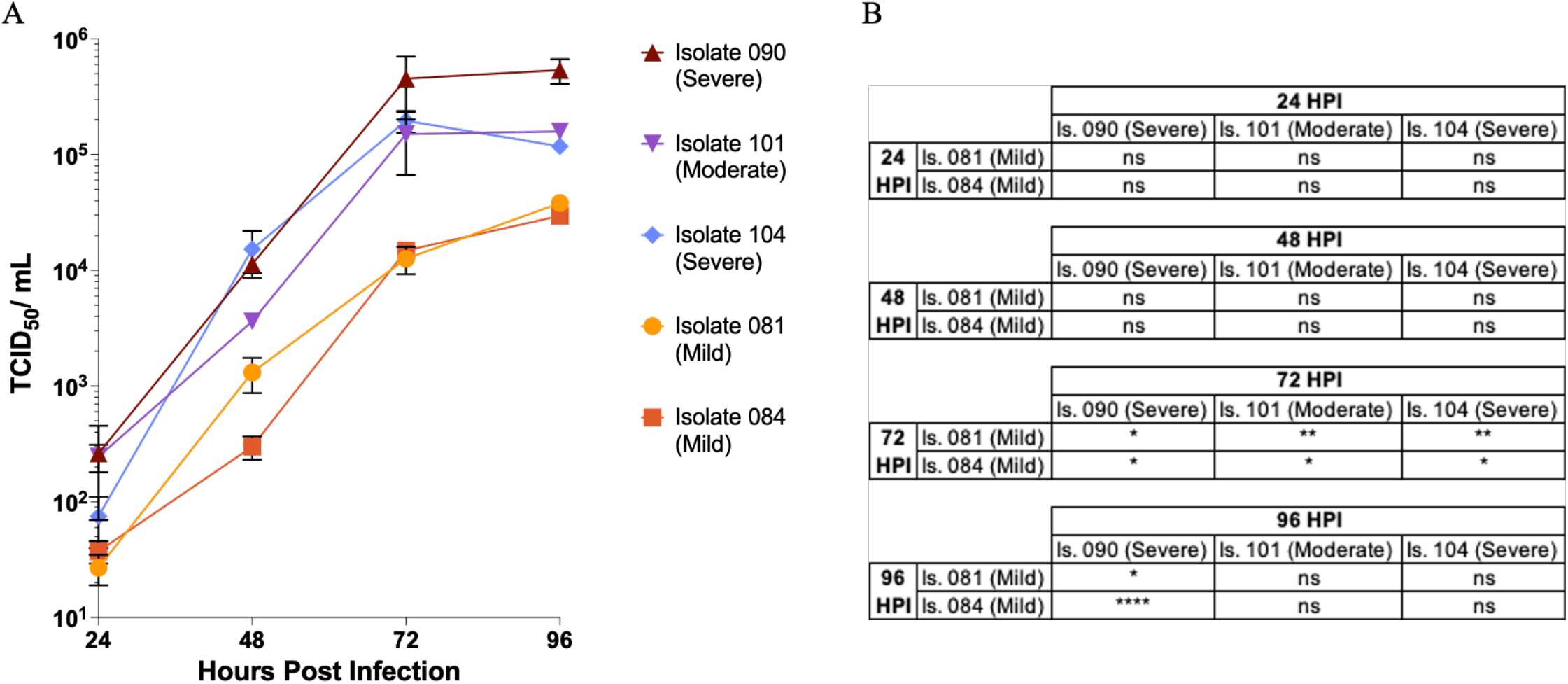
Temporal analysis of RSV isolate replication **A)** A549 cells were infected with RSV isolates at an MOI of 0.01. Extracellular RSV was isolated, and viral titer was determined by TCID_50._ Samples were collected at 24H intervals over 96 hours. Data shown represent the mean of duplicate measurements +/-SD **B)** Table summarizing the statistical significance of differences in viral titer between mild, moderate, and severe strains at each timepoint. ns (not significant), * (p < 0.05), ** (p < 0.01), *** (p < 0.001), **** (p < 0.0001). Abbreviations: TCID_50_, 50% tissue culture infectious dose; MOI, multiplicity of infection; Is, Isolate; HPI, Hours post-infection.

Given the evidence that isolates from moderate/severe RSV disease replicate to higher titers than those from mild cases (**Figure 2)**, we employed predictive protein modeling to elucidate the structural consequences of associated mutations. Structural modeling suggested that these mutations, particularly in the L and G proteins, may enhance viral fitness by altering protein stability, dimerization, and host interaction. All isolates encoded a 321-aa G protein. N-glycosylation prediction showed a stepwise increase with severity: mild isolates had two sites, moderate/severe had three to four (**S1 Figure**). A previous report demonstrated that glycosylation of the C-terminal domain of the G protein reduced reactivity with specific antibodies, suggesting that various glycosylation patterns of RSV may contribute to evasion of the host’s humoral immune response (26).

L polymerase sequences revealed five amino acid differences between mild and moderate/severe isolates: two in the RdRp domain (143, 179) and three within the connector region (1653, 1661, 1715) (**Table 2**). Structural modeling based on Protein Data Bank 6UEN (27) highlighted these substitutions but limited interpretation to the experimentally resolved RdRp and capping regions (**S2 Figure**). An additional aspartate residue in the finger domain of the RSV L protein, specifically at position AA143, may have significant implications for polymerase function. Introduction of an additional aspartate in this region could alter the electrostatics of the template entry pore or influence RNA binding affinity (28,29). Such changes may impact processivity or regulation of RNA synthesis, thereby affecting viral replication efficiency and, consequently, clinical outcomes.

Reported associations between RSV viral load and disease severity in young children have been inconsistent, likely due to variations in study design, analytical methods, host immune responses, and intrinsic viral factors. Although our sample size is small, the compelling molecular findings presented here highlight the potential for specific pathoadaptive mutations to influence evolving RSV virulence and clinical outcomes. These results may help to explain previously reported inconsistencies and highlight the need for larger studies, reverse genetics, and in vivo models to elucidate the specific mechanisms by which RSV phenotype shapes disease severity.

## Acknowledgments

We would like to express our immense gratitude to our colleagues Carlie DeFelice, Dr. Kelly Hawley, Judy Kalinowski, Carson Karanian, Stephanie Lesmes, Dr. Bo Reese, and Jordan Wolf for their vital contribution to the project, including participant recruitment, biospecimen collection, processing and testing, and data capture.

## Supplemental Information

**S1 Figure.**
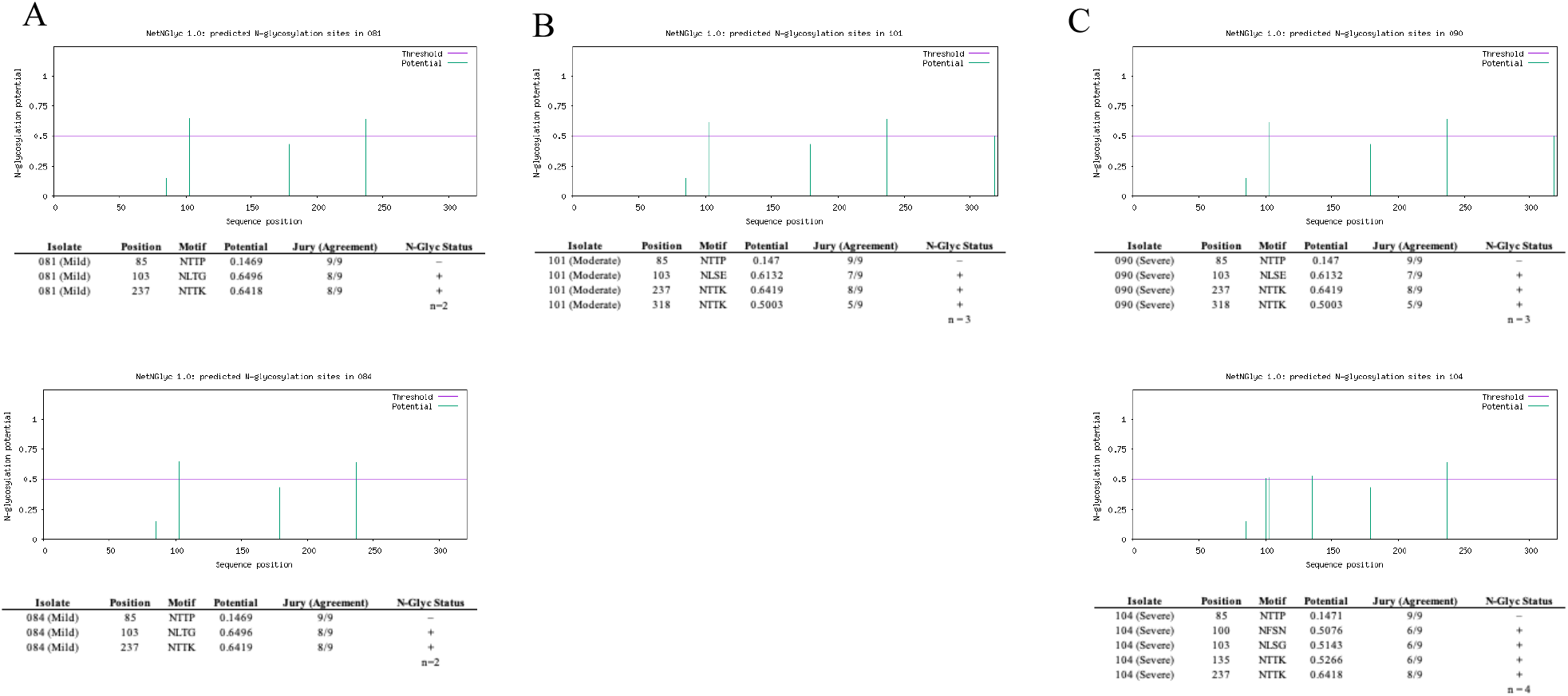
Predicted N-glycosylation sites in RSV Attachment G protein A) in isolates from cases of mild disease (Is. 081, Is. 084), B) in isolates from moderate cases of disease (Is. 101) and C) in isolates from severe cases of disease (Is. 090, Is. 104). N-glycosylation potential threshold = 0.5 (purple line). Potential is the predicted likelihood of glycosylation (green). Jury is the number of prediction tools in agreement out of 9.

**S2 Figure.**
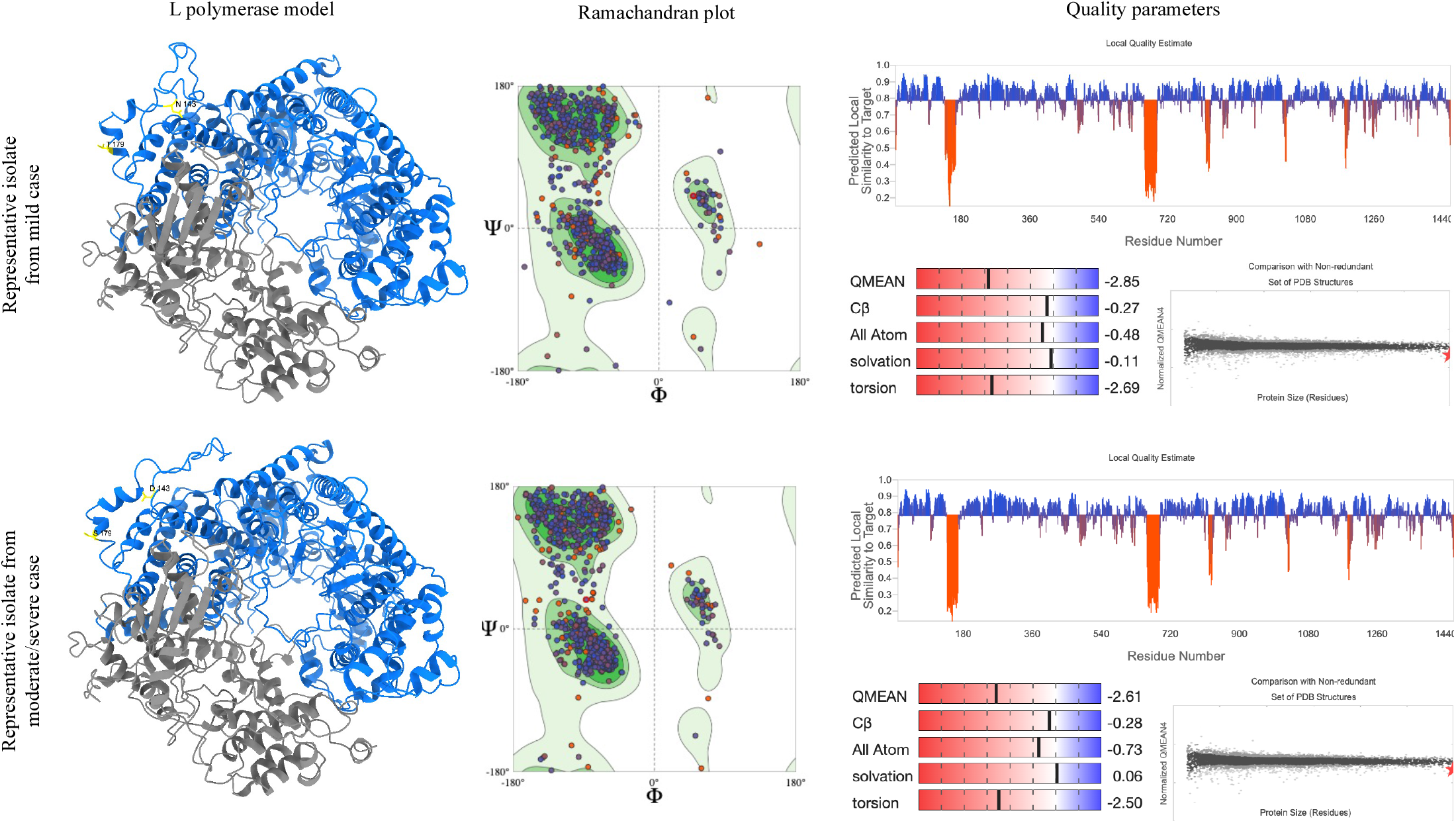
Structural model of RSV L proteins of representative isolates from mild and moderate/severe cases using homology modelling based on the RSV polymerase complex structure (PDB: 6UEN). Ribbon representation of L proteins are shown on left (yellow, pathoadaptive mutations; blue, RNA-dependent RNA polymerase domain; grey, PRNTase capping domain). Ramachandran plots are shown for each model (center) followed by quality parameters of each model. T, Threonine; N, Asparagine; S, Serine; D, Aspartate;

**S1 Table.** Clinical Disease Severity Score (CDSS)

|  | **0** | **1** | **2** | **3** |
| --- | --- | --- | --- | --- |
| **O<sub>2</sub> Saturation** | >95% in RA | 92%-95% in RA | <92% in RA or<br>suppl O<sub>2</sub> ≤35% | Intubated/CPAP or<br>suppl O<sub>2</sub>>35% |
| **Respiratory Rate** | ≤6 m <40/min<br>>6 m <30/min | ≤6 m 41-55/min<br>>6 m 31-45/min | ≤6 m 56-70<br>>6 m 46-60 | ≤6 m >70<br>>6 m >60 |
| **Retractions** | No | Subcostal + | Subcostal ++ or<br>intercostal | Subcostal +++ or<br>intercostal and<br>suprasternal |
| **Auscultation** | Normal | Crackles/<br>ronchi | Any wheezing | Exp/insp wheezing<br>without stethoscope<br>or no breath sounds<br>heard |
| **Activity** | Normal, alert | Abnormal, fussy, or<br>decreased oral<br>intake | Unable to maintain<br>hydration (IV fluids<br>or NG feeds) | Intubated |

<sup>a</sup>Score from 0-15: Mild (0-5), Moderate (6-9), Severe (≥10).
<sup>b</sup>CPAP, continuous positive airway pressure; exp, expiratory; insp, inspiratory; IV, intravenous; m, months; min, minute; NG, nasogastric tube; O<sub>2</sub>, oxygen; RA, room air; suppl, supplemental; +, mild; ++, moderate; +++, severe.

